# 15-PGDH Inhibition Overcomes Muscle Regenerative Deficit Seen With GLP1-Receptor Agonist–Induced Weight Loss

**DOI:** 10.64898/2026.02.26.708119

**Authors:** Minas Nalbandian, Jameel Lone, Emmeran Le Moal, Ireh Kim, Yutong Kelly Li, Peggy Kraft, Meng Zhao, Kassie Kolacar, Zeyuan Zhang, Katrin J. Svensson, Helen M. Blau

## Abstract

Glucagon-like peptide-1 receptor agonists (GLP-1 RAs), including long-acting semaglutide, are revolutionary anti-obesity therapies. However, emerging evidence indicates that weight loss may come at the expense of skeletal muscle mass, a tissue essential for mobility, metabolic regulation, and overall health. Here, we show that an inhibitor of the gerozyme 15-hydroxyprostaglandin dehydrogenase (PGDHi), which boosts PGE2 levels, increases skeletal muscle mass, strength, and regeneration in the presence of semaglutide. We find that in a high fat diet-induced mouse model of obesity, semaglutide alone induces significant loss of muscle mass, while retaining contractile function. However, muscle regeneration and recovery of strength post-injury are hindered by semaglutide. This regenerative deficit is due to impeded stem cell function, which is overcome if mice are treated with a combination of PGDHi and semaglutide. Our data show that GLP-1–mediated weight loss interferes with this key muscle-building function, which PGDHi co-treatment counteracts to promote proper muscle regeneration and restored strength.

## Introduction

Obesity is associated with reduced life expectancy and quality of life (1–3). It is a major health challenge and contributor to the rising incidence of type 2 diabetes, cardiovascular disease, nonalcoholic fatty liver disease, and several forms of cancer. Despite sustained public-health efforts, the global prevalence of obesity has more than doubled since the 1980s, rising from approximately 6% to over 14% in adults, and it is escalating worldwide, underscoring the urgent need for effective and scalable therapeutic strategies (4). While lifestyle interventions are fundamental, pharmacological therapies that achieve sustained weight loss are critical for managing obesity and its numerous comorbidities. Among the most effective agents currently available are glucagon-like peptide-1 receptor agonists (GLP-1 RAs), such as semaglutide, which mimic endogenous incretin signaling to regulate glucose metabolism (5). These drugs act in the central nervous system to reduce appetite, and peripherally to delay gastric emptying and improve insulin secretion and sensitivity (6).

Large randomized clinical trials consistently demonstrate that GLP-1 RAs induce significant and durable reductions in body weight (7–9). For example, six weeks of weekly semaglutide injections resulted in an average 16% body weight reduction in obese adults (7), a degree of efficacy that rivals bariatric surgery (10). Similar weight loss benefits have been reported in obese adolescents, evidence of the clinical relevance of these agents to younger populations (11). The success of GLP-1 RAs has led to their widespread adoption for multiple indications. In addition to their metabolic effects, these agents offer cardioprotective (12) and renal benefits (13), which explains their growing appeal in treating obesity as a systemic disease.

A major caveat is that the weight loss associated with semaglutide results not just from the loss of fat mass, but also from the loss of lean mass including muscle (14, 15). Large-scale randomized clinical trials have shown that GLP-1 RA–induced weight loss impacts both fat and lean tissue, with adults losing over 5 kg of lean mass after 68 weeks of treatment (7). In preclinical models, semaglutide has been associated with significant muscle atrophy, further reinforcing concerns about its impact on skeletal muscle mass and integrity (16, 17). This is clinically relevant, as the loss of skeletal muscle can compromise strength, metabolic health, and quality of life. Preserving muscle during weight loss has long been a therapeutic objective.

Relatively little is known about how semaglutide affects muscle function, particularly in the context of tissue remodeling as occurs after injury, surgery, and exercise. These processes rely on cycles of minor damage and subsequent regeneration to promote gains in muscle mass and strength (18). An understanding of the impact of GLP-1RA treatments on regenerative capacity in obese individuals is particularly important, as it may dictate their response to physical activity, rehabilitation post-surgery, and overall effectiveness of exercise-based interventions.

We previously identified the enzyme 15-hydroxyprostaglandin dehydrogenase (15-PGDH), as a gerozyme, a molecular driver of muscle wasting post-injury and in aging (19–21). 15-PGDH degrades prostaglandin E2 (PGE2), a lipid metabolite that has pleiotropic beneficial effects on skeletal muscle. PGE2 promotes muscle stem cell proliferation and tissue regeneration, myofiber mitochondrial number and function, and restores neuromuscular connectivity (19–22). We showed that pharmacological or genetic inhibition of 15-PGDH (PGDHi) enhances PGE2 signaling and increases muscle mass and strength, especially in aged mice (19, 21).

Here we test the effects of semaglutide, alone and in combination with PGDHi, on skeletal muscle size, strength, and regeneration in young adult obese mice. Our findings reveal that semaglutide reduces muscle mass but preserves force under steady-state conditions. However, semaglutide induces muscle stem cell dysfunction resulting in impaired regeneration and smaller myofibers. Treatment with PGDHi overcomes this deficit by boosting muscle stem cell proliferation, providing the building blocks for muscle tissue repair and restoring myofiber size, without interfering with semaglutide’s metabolic benefits. Our results identify 15-PGDH as a critical node in muscle adaptation to metabolic therapies. Further, they provide support for PGDHi as a therapeutic strategy to enable muscle growth and acquisition of strength during pharmacologically induced weight loss.

## Results

### Semaglutide triggers a loss of muscle mass but preserves muscle function

Glucagon-like peptide-1 receptor agonists (GLP-1 RAs) result in significant body weight loss, however, a portion of this weight loss results from a reduction in lean body mass, particularly skeletal muscle(17, 23, 24), which is a concern given the critical role of skeletal muscle in systemic metabolism and movement. Thus, there is a major unmet need for drugs that preserve muscle in conjunction with GLP-1 RA interventions. While numerous GLP-1RA studies have assessed loss of muscle mass, few have measured muscle function in terms of strength. We reasoned that treatment with an inhibitor of 15-hydroxyprostaglandin dehydrogenase (15-PGDH) could overcome the loss of muscle mass and function observed with GLP-1 RAs. 15-PGDH degrades prostaglandin E2 (PGE2) and is a gerozyme that increases with injury and aging (21, 25, 26). To test if an inhibitor of 15-PGDH (PGDHi) could preserve muscle mass and function during semaglutide-induced weight loss, we compared the effects of semaglutide, a GLP-1 RA approved for clinical use under the brand name Ozempic or Wegovy, alone and in combination with PGDHi on body composition and skeletal muscle mass and function in a mouse model of high fat diet-induced obesity (HFD).

To induce obesity, young adult (8 wk of age) male C57BL/6J mice were fed a high-fat diet (HFD; 60% kcal from fat) for 12 weeks. Animals were then randomized by body weight and baseline plantar flexion maximal tetanic force into four treatment groups: vehicle control (Veh), PGDHi, semaglutide, and combined (PGDHi + semaglutide) (**Fig. 1A**). Body weight was recorded daily throughout the intervention, and maximal strength was assessed in vivo as plantar flexion torque at baseline, three weeks, and five weeks after initiation of treatment. In agreement with reports by others (17, 27), semaglutide treatment led to a sustained and significant reduction in body weight, reaching ∼25% total body weight loss by week five (**Fig. 1B–C**). Importantly, co-administration with PGDHi did not significantly attenuate the weight-lowering effect of semaglutide, indicating that PGDHi alone does not induce weight loss or interfere with the appetite-suppressing effect of semaglutide. Semaglutide also significantly reduced heart weight (**Supplementary Fig. 1A, B**) and improved glucose tolerance (**Supplementary Fig. 2A, B**) In agreement with reports by others (28–30). Notably, PGDHi alone had no detectable effect on either outcome. To further assess changes in adiposity, we isolated and weighed major adipose depots: inguinal (iWAT), epididymal (eWAT), and brown adipose tissue (BAT). Both semaglutide and the PGDHi + semaglutide significantly reduced iWAT, eWAT, and BAT weights compared to PGDHi alone or vehicle controls (**Fig. 1D**), in agreement with histological analyses (**Fig. 1E**). These data confirm that semaglutide effectively reduces fat mass and that PGDHi does not add to or interfere with this process.

**Figure 1.**
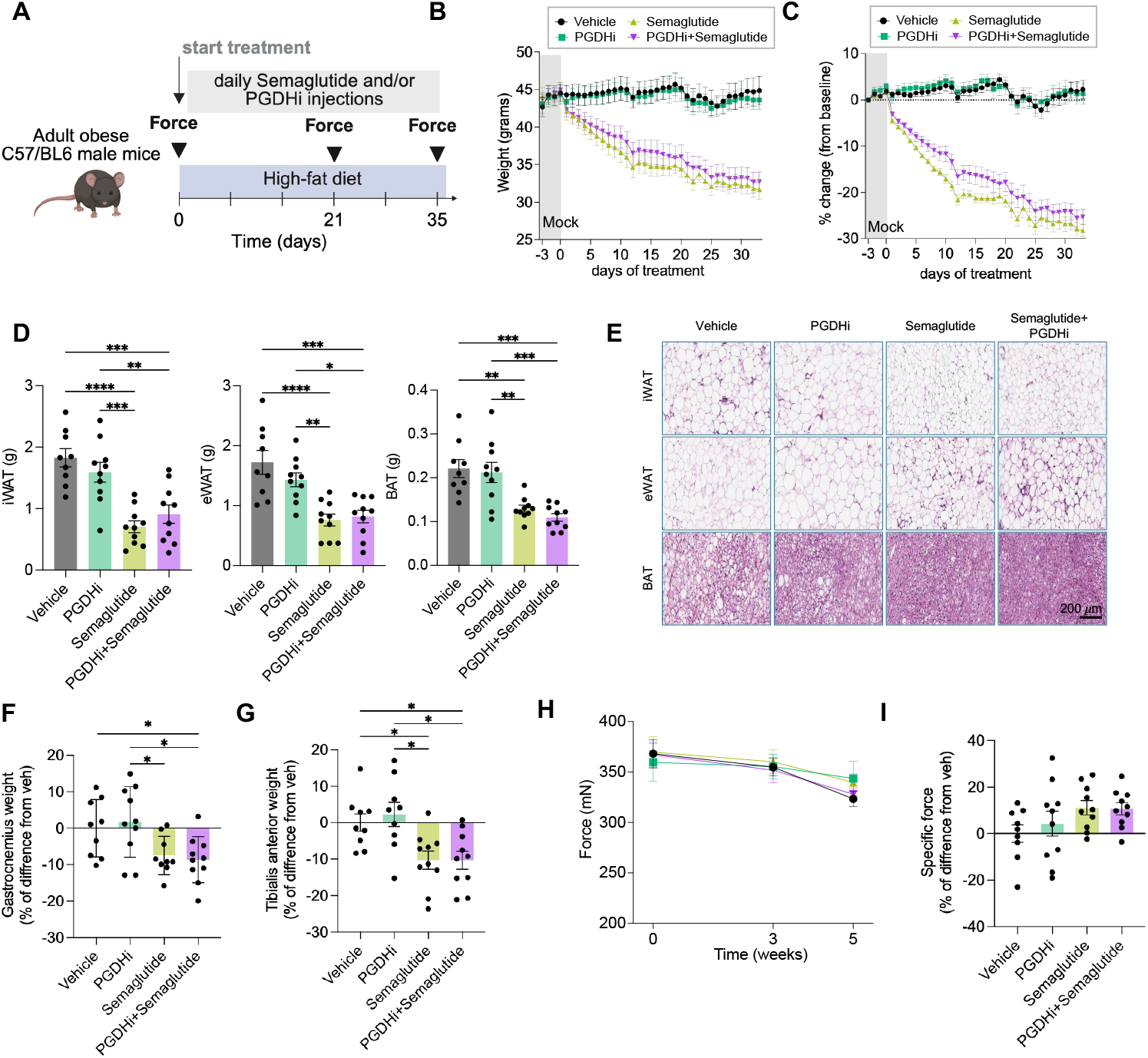
Semaglutide causes body weight and muscle loss while preserving muscle function in obese mice. (A) Experimental schematic. 8 weeks old mice were fed with High Fat Diet (HFD) for 12 weeks and then treated with semaglutide and/or PGDHi for 5 weeks while maintaining the HFD. Maximal tetanic force was measured at the start of the treatment, at 3- and 5-weeks post treatment. (B) Body weight in grams. (C) Proportion of body weight change relative to baseline. (D) Wet tissue weights for iWAT (inguinal adipose tissue), eWAT (epididymal adipose tissue), and BAT (brown adipose tissue). (E) Representative H&E staining images of iWAT (inguinal adipose tissue), eWAT (epididymal adipose tissue), and BAT (brown adipose tissue). (F-G) Relative wet tissue weight of Gastrocnemius and Tibialis anterior muscles. (H) Maximal tetanic force across the treatment. (I) Relative specific force at the end of the treatment (5 weeks). n=8-10 animals per group. Data are presented as mean ± S.E.M. Statistic significance was analyzed by two-way ANOVA for multiple comparisons. * p < 0.05; ** p < 0.01; *** p < 0.001; **** p < 0.0001. ### p < 0.001 for Vehicle vs Semaglutide and Vehicle vs PGDHi + Semaglutide.

To determine if the marked loss of body weight was accompanied by a loss of skeletal muscle mass and strength, we weighed the gastrocnemius (GA) and tibialis anterior (TA) muscles of obese mice and found that semaglutide-treated mice showed a significant reduction in muscle mass (**Fig. 1F–G**). However, histological analyses of gastrocnemius sections revealed no detectable differences in myofiber size across treatment groups **(Supplementary Fig. 3A, B)**, suggesting that the observed changes in muscle mass were below the threshold needed to produce measurable alterations in fiber size, as previously reported (31). Similarly, muscle strength remained unchanged across all groups throughout the intervention (**Fig. 1H**). To evaluate muscle quality, we calculated specific force (force normalized to muscle mass) and found that with semaglutide, either alone or together with PGDHi, did not increase specific force (**Fig. 1I**), Together, these findings suggest that semaglutide induces weight loss predominantly through fat reduction, but also reduces skeletal muscle mass, consistent with prior mouse and human data (7, 17, 32). Importantly, PGDHi does not interfere with semaglutide-induced loss of weight.

### 15-PGDH inhibition significantly increases recovery of muscle strength following Injury in semaglutide-treated obese mice

We sought to determine if GLP-1 RA treatment would alter the regenerative capacity of muscle. Skeletal muscle possesses remarkable plasticity and can readily adapt to a range of external stimuli such as exercise, metabolic stress or tissue injury (33–36). Skeletal muscle relies on a resident muscle stem cell (MuSC) pool to rebuild muscle tissue following injury. Regeneration is a highly orchestrated process requires multiple rounds of MuSC proliferation with high energetic demands that would be difficult to achieve under caloric restriction such as occurs with GLP1-RA therapy. To test if regenerative capacity was altered, young adult obese mice were treated for three weeks with vehicle, semaglutide, PGDHi, or the drug combination. At the end of the third week, gastrocnemius muscles were injured via intramuscular injection of notexin. Semaglutide and PGDHi treatments were continued for an additional two weeks post-injury (**Fig. 2A**) at which time force was assessed and gastrocnemius muscles were harvested and weighed to assess muscle mass. As observed in the absence of injury, semaglutide significantly reduced gastrocnemius muscle weight compared to vehicle-treated controls, whereas PGDHi alone did not alter muscle mass (**Fig. 2B**). Notably, despite the reduction in muscle mass, force measurements revealed a significant increase in muscle performance in mice co-treated with semaglutide and PGDHi, not seen with either treatment alone (**Fig. 2C–D**). Maximal strength, assessed two weeks post-injury, was significantly higher in the PGDHi + semaglutide group compared to all other groups (**Fig. 2C–D**). Additionally, specific force was markedly increased in the co-treated mice, suggesting that the increase in efficiency of regeneration resulted in enhanced muscle quality (**Fig. 2E**). These findings suggest that PGDHi augments muscle functional recovery post-injury in semaglutide-treated mice.

**Figure 2.**
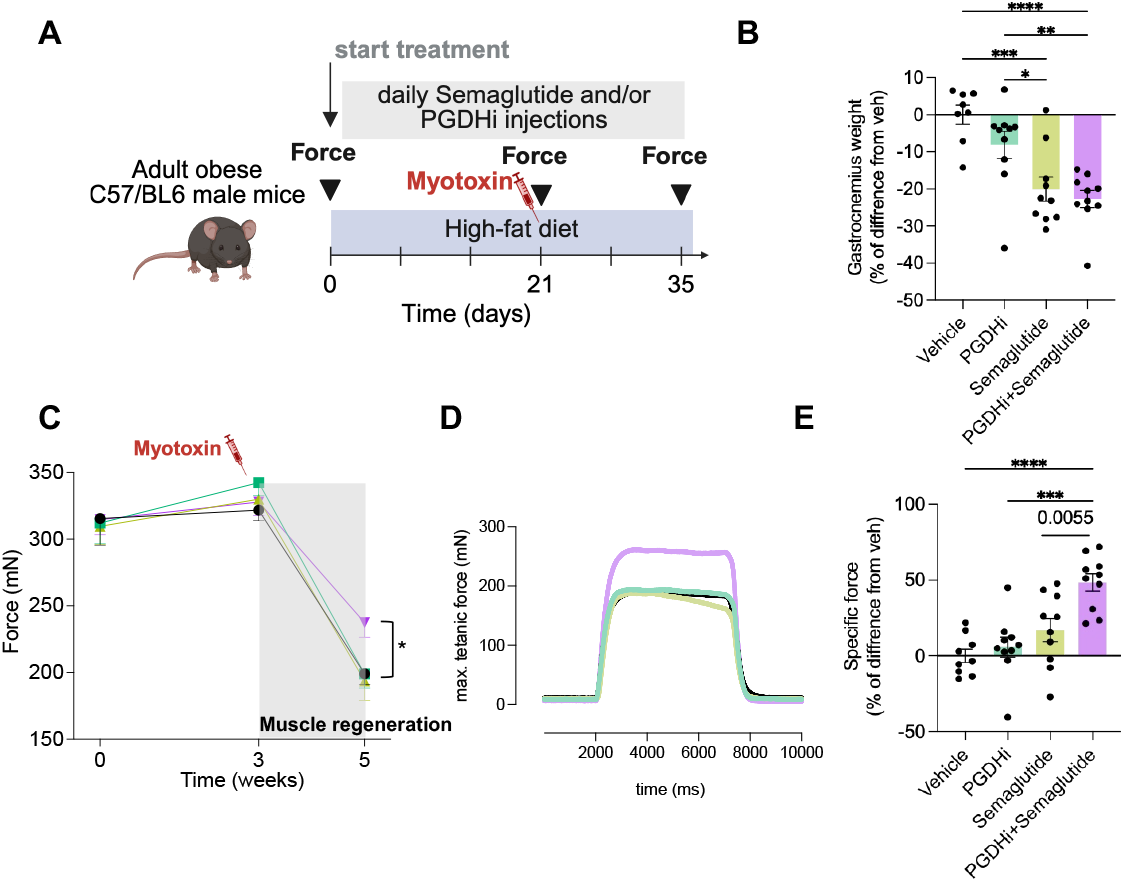
Co-administration of semaglutide and PGDHi increases muscle force 2 weeks post injury in obese mice. (A) Experimental schematic. Mice were fed with High Fat Diet (HFD) for 12 weeks and then treated with semaglutide and/or PGDHi for 5 weeks while maintaining the HFD. At 3 weeks of starting the treatment, the left gastrocnemius muscle were injured with a notexin injection. Maximal tetanic force was measured at the start of the treatment, at 3-and 5-weeks post treatment. (B) Gastrocnemius wet tissue weight. (C) Maximal tetanic force across the treatment. P values in the plot are for comparisons with PGDHi+sema group, all shown comparison were P <0.05. One asterisk denotes that all pairwise comparisons vs PGDHi+sema were significant. (D) Representative traces of the maximal tetanic force at week 5. (E) Relative specific force at the end of the treatment (5 weeks). n=8-10 animals per group. Data are presented as mean ± S.E.M. Statistic significance was analyzed by two-way ANOVA for multiple comparisons. * p < 0.05; ** p < 0.01; *** p < 0.001; **** p < 0.0001.

To elucidate potential structural correlates of this improved function, we performed histological analyses on gastrocnemius cross-sections collected two weeks post-injury. Tissue sections were immunostained for laminin to delineate fiber boundaries, as well as hematoxylin and eosin (H&E) and Sirius Red staining to assess overall morphology and fibrosis, respectively (**Fig. 3A**). Quantification of regenerating fibers, identified by centrally located nuclei, revealed that semaglutide-treated muscles displayed significantly smaller fiber diameters compared to all other groups (**Fig. 3B; Supplementary Fig. 4**), indicating impaired myofiber growth during regeneration. Notably, this reduction in fiber size was fully rescued in mice co-treated with semaglutide and PGDHi.

**Figure 3.**
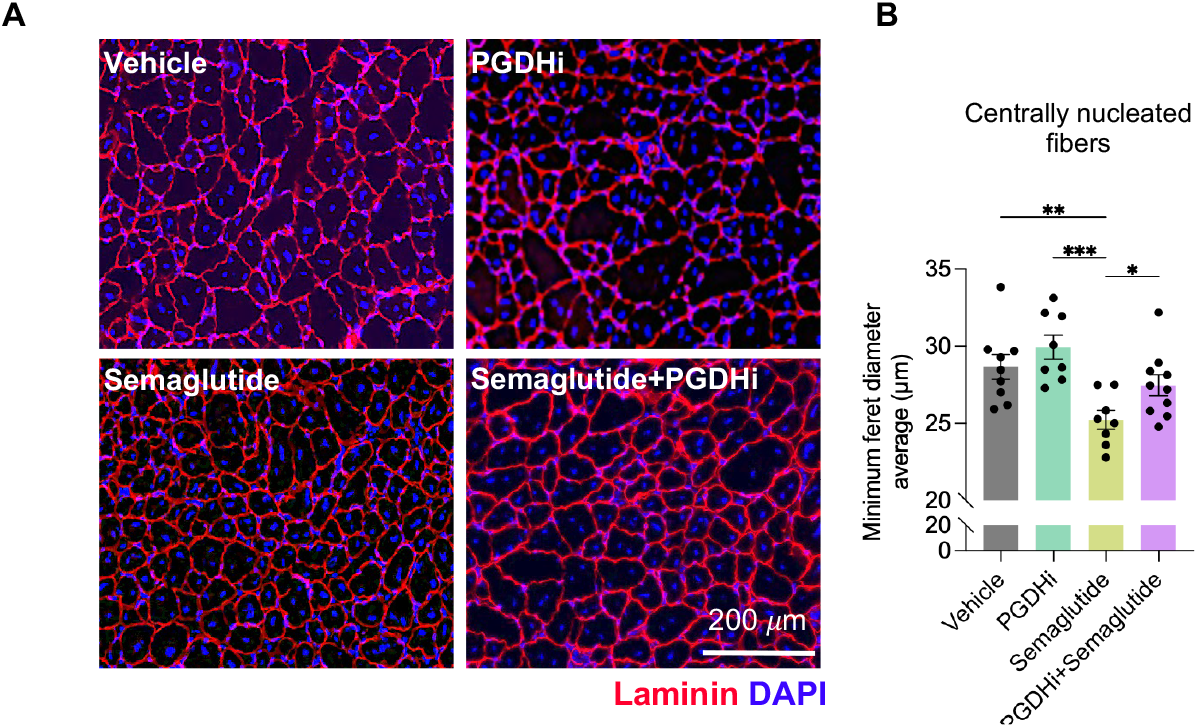
PGDHi restores myofiber size blunted by semaglutide in regenerating muscle. (A) Representative images of gastrocnemius muscles at 2 weeks post injury. Laminin (red) and DAPI (blue) staining. (B) Quantification of the minimum Feret diameter of centrally nucleated fibers from the laminin staining. n=8-10 animals per group. Data are presented as mean ± S.E.M. Statistic significance was analyzed by two-way ANOVA for multiple comparisons. * p < 0.05; ** p < 0.01; *** p < 0.001.

Sirius Red and H&E staining revealed regions of fibrotic tissue and presumptive calcification, which were confirmed by Alizarin red staining as calcium deposits in the gastrocnemius of vehicle-treated obese mice **(Supplementary Fig. 5A–C**). These aggregates were specific to regenerating obese muscles. These pathological features persisted with PGDHi but were reduced in the presence of GLP1-RA with and without PGDHi. These results suggest that GLP-1 receptor agonism mitigates the fibrotic and degenerative remodeling typically observed during muscle regeneration in obese mice (37). Notably, the accumulation of calcium deposits is similar to what has been reported for DMD muscle where it has been linked to a regenerative defect (38–40). Taken together, these data indicate that semaglutide has a dual beneficial effect during muscle regeneration: it protects against fibrotic and calcific tissue degeneration. Importantly, PGDHi co-administration ameliorates the semaglutide-associated reduction in fiber size while preserving the anti-fibrotic and anti-calcification effects in obese metabolically compromised mice. These findings show that PGDHi and GLP-1 have synergistic effects on muscle regeneration and acquisition of strength.

### 15-PGDHi enhances muscle stem cell proliferation and regeneration in semaglutide-treated obese mice

We reasoned that the relatively diminished CSA of muscle fibers post injury in semaglutide-treated obese mice results from compromised regenerative capacity. Muscle regeneration relies on the activation, proliferation, and differentiation of resident muscle stem cells (MuSCs), also known as satellite cells (41, 42). We have previously demonstrated that prostaglandin PGE2 is essential to MuSC function. In mice in which the receptor for PGE2 EP4 is genetically ablated, regeneration is severely blunted and strength is not restored post-injury (22). Similar results are observed if endogenous PGE2 synthesis is abrogated by treatment to mice post injury with an NSAID. This failure to regenerate and restore muscle function after injury is due to a requirement for PGE2 for MuSC proliferation and survival. In the absence of this signaling pathway they cannot meet the needs of expansion to repair the muscle damage (22).

To test if pharmacological inhibition of 15-PGDH enhances MuSC proliferation in injured muscles, gastrocnemius muscles of obese mice were injured via intramuscular notexin injection, and drug treatments were continued for five days. To track *in vivo* MuSC proliferation, mice received continuous supplementation of 5-ethynyl-2’-deoxyuridine (EdU) in the drinking water starting from the day of injury. Based on established kinetics of the MuSC proliferative response, which peaks around 5 days post-injury (43, 44), muscles were harvested at this time point for histological and molecular analyses (**Fig. 4A**). As expected from previous experiments, semaglutide-treated animals (both semaglutide and semaglutide/PGDHi groups) exhibited significant reductions in total body weight, heart weight, and fat mass compared to vehicle controls (**Supplementary Fig. 6A–E**). The regenerating gastrocnemius muscle mass was also significantly decreased in semaglutide-treated mice, regardless of PGDHi co-administration (**Fig. 4B**). These data indicate that semaglutide maintains its systemic metabolic effects under injury conditions, including weight loss and reductions in fat, cardiac and skeletal muscle tissue mass.

**Figure 4.**
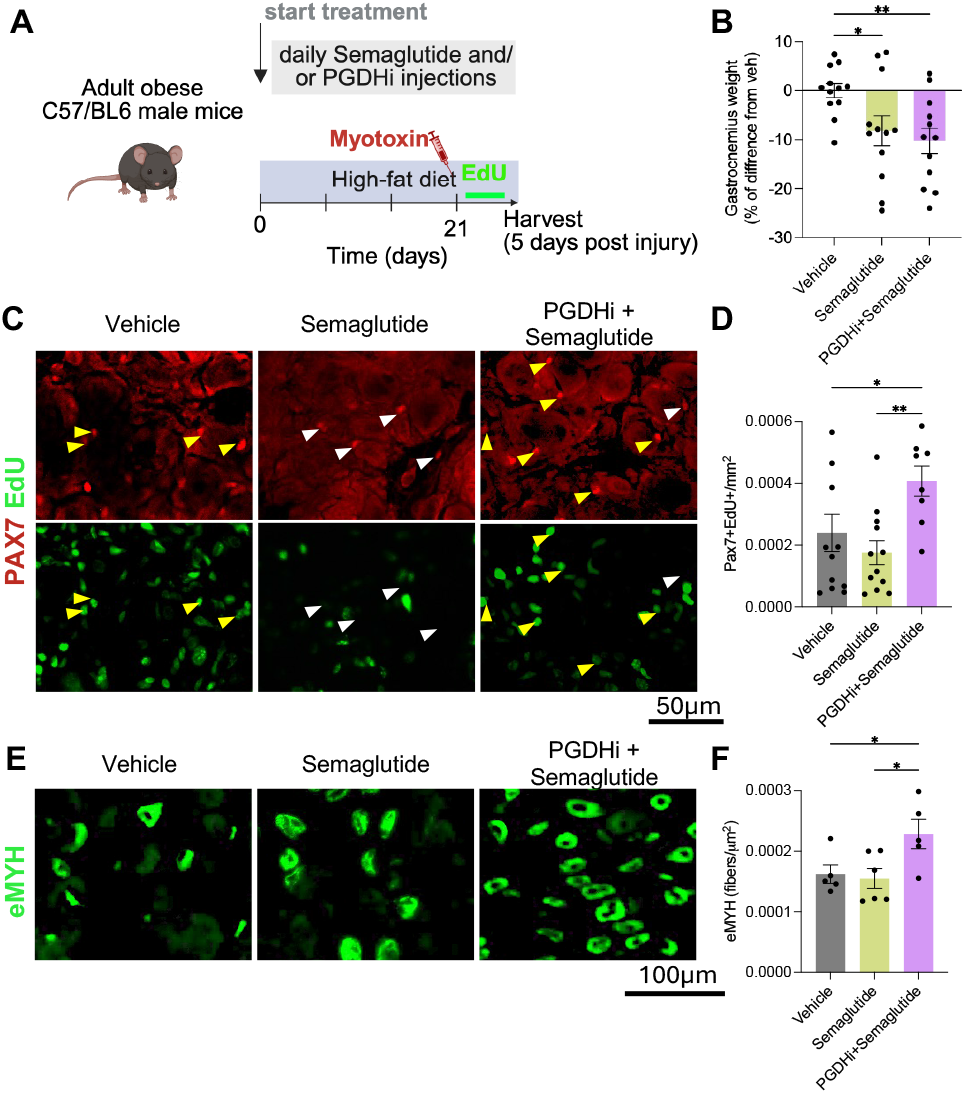
Co-administration of semaglutide and PGDHi enhances stem cell function in injured obese mice. (A) Experimental schematic. Mice were fed with High Fat Diet (HFD) for 12 weeks and then co-treated with semaglutide and PGDHi (or the respective controls) for 3 weeks while maintaining the HFD. At 3 weeks of starting the treatment, myotexin was used to induce injury in the left gastrocnemius muscle and EdU supplementation initiated. Sema and PGHDI treatment continued throughout this period. Mice were sacrificed and tissue collected 5 days after injury. (B) Relative gastrocnemius wet tissue weight. (C) Representative gastrocnemius cross sectional areas at day 5 post injury stained for PAX7 (red), EdU (green), Laminin (white), and DAPI (blue). Yellow triangles indicate PAX7^+^/EdU^+^ cells; White triangles indicate PAX7^+^/EdU^-^ cells. (D) Quantification of PAX7^+^/EdU^+^ cells per area. (E) Representative gastrocnemius cross sectional areas at day 5 post injury stained for eMYH (green), Dystrophin (white), EdU (Red), and DAPI (blue). (F) Quantification of eMYH^+^ fibers per area. (B,D) n=6 animals per group, right and left gastrocnemius were injured and included in the analysis. (F) n=5-6 animals per group, only right gastrocnemius were used. Data are presented as mean ± S.E.M. Statistic significance was analyzed by two-way ANOVA for multiple comparisons. * p < 0.05; ** p < 0.01; *** p < 0.001; **** p < 0.0001.

To determine if PGDHi enhances MuSC proliferation during regeneration, we performed immunofluorescence staining on cryosections of injured gastrocnemius muscles at five days post-injury. Sections were co-labeled for EdU, indicative of DNA synthesis, and PAX7, the canonical marker for quiescent and activated MuSCs. This dual labeling enabled identification of actively proliferating MuSCs (EdU^+^/PAX7^+^) (**Fig. 4C**). Quantification revealed a significant increase in the number of EdU^+^/PAX7^+^ cells per section in the PGDHi + semaglutide group compared to semaglutide alone or vehicle controls (**Fig. 4D**). These results indicate that co-treatment with semaglutide and PGDHi significantly augments MuSC proliferation, as evidenced by increased numbers of EdU^+^/PAX7^+^ cells, compared to either semaglutide or vehicle alone. These findings suggest that 15-PGDH inhibition overcomes the deficit in MuSC proliferation seen with semaglutide-induced weight loss.

To assess the downstream consequences of enhanced MuSC proliferation on myofiber regeneration, we analyzed the expression of embryonic myosin heavy chain (eMyHC), a marker of nascent regenerating myofibers. Immunostaining revealed a greater number of eMyHC-positive fibers in PGDHi + semaglutide treated muscles compared to vehicle or semaglutide only treated groups (**Fig. 4E**), suggestive of more robust or accelerated muscle regeneration in PGDHi treated mice relative to those treated only with semaglutide.

To further dissect the mechanism by which PGE2 promotes MuSC proliferation, we isolated α7-integrin^+^ MuSCs from uninjured 12-week-old C57BL/6 mice. To mimic the nutrient deficit generated by semaglutide, we cultured the MuSCs under nutrient-depleted conditions, and supplemented the media with PGE2, EP2/EP4 receptor inhibitors (EPi) to block PGE2 binding to its receptors, or a CREB phosphorylation inhibitor (pCREBi) (**Fig. 5A)**. CREB was targeted because activation of EP2/EP4 receptors is known to increase intracellular cAMP and stimulate CREB phosphorylation, a key transcriptional mediator of PGE2-driven proliferation and survival (20, 22). Thus, EPi tests whether the proliferative effect of PGE2 is receptor-dependent, whereas pCREBi tests whether downstream cAMP–CREB signaling is required. EdU incorporation assays revealed that PGE2 treatment significantly increased MuSC proliferation compared to vehicle control, whereas co-treatment with either an EP2 or EP4 inhibitor or a pCREBi attenuated this effect (**Fig. 5B–C**). These results indicate that the proliferative effects of PGE2 on MuSCs require EP receptor signaling and downstream CREB activation, providing a link between 15-PGDH inhibition, PGE2, and MuSC proliferation.

**Figure 5.**
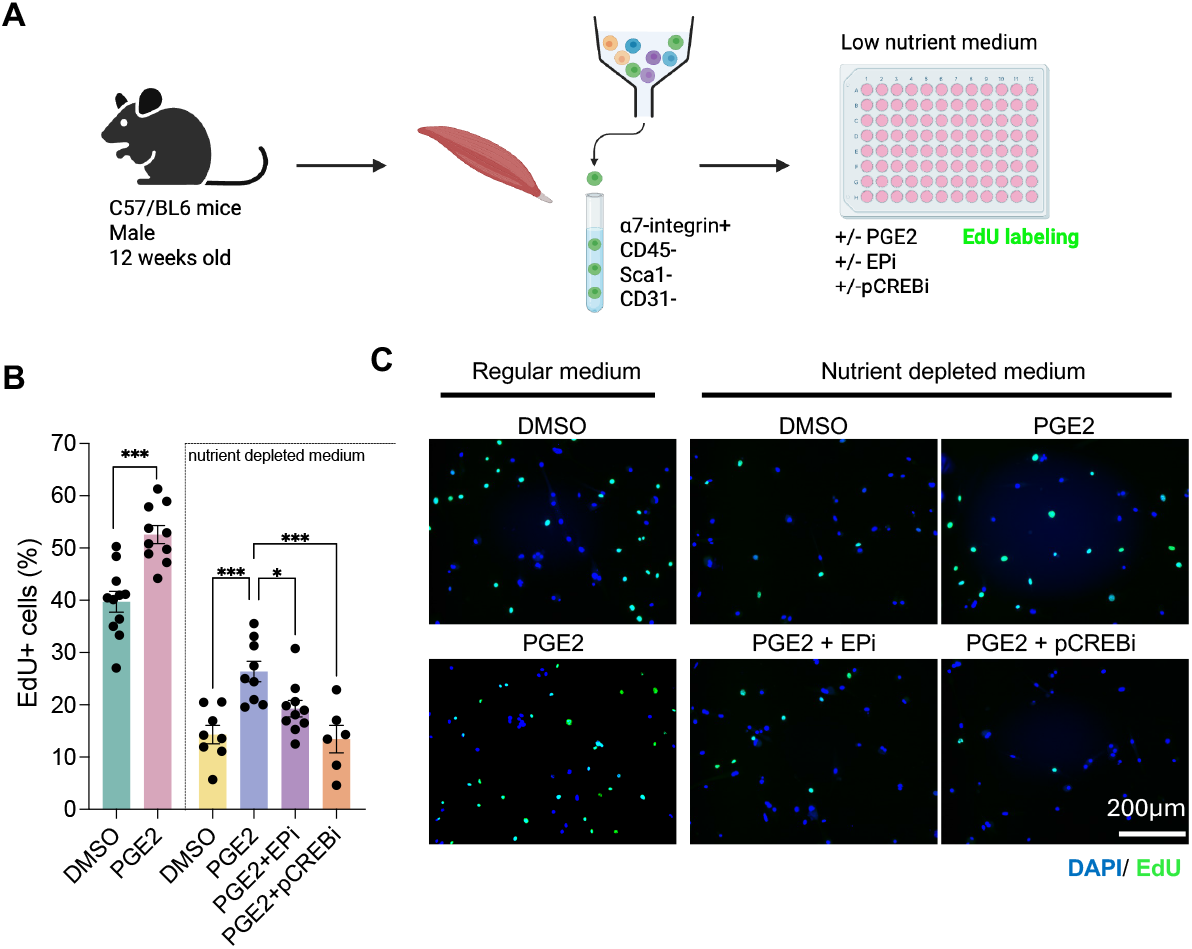
PGE2 enhances MuSC proliferation in vitro in low nutrient conditions. (A) Schematic of experimental design. MuSCs (α7-integrin^+^, CD45^−^, Sca1^−^, CD31^−^) were isolated from skeletal muscle of 12-week-old male C57BL/6 mice and cultured in regular or nutrient-depleted medium with the indicated treatments. EdU incorporation was used to assess proliferation. (B) Quantification of EdU^+^ MuSCs in nutrient-depleted medium. PGE2 increased proliferation compared with DMSO, while co-treatment with EPi (EP2/EP4 inhibitor) or pCREBi (CREB inhibitor) reduced this effect. Data are presented as mean ± SEM. Statistical significance: * p < 0.05; ** p < 0.01; *** p < 0.001; **** p < 0.0001. (C) Representative immunofluorescence images of EdU (green) and DAPI (blue) under the indicated conditions. Scale bar, 200 μm.

Together, these data show that while semaglutide reduces muscle calcification, it fails to promote effective muscle regeneration in diet-induced obesity mice. In contrast, concurrent GLP-1 receptor agonist and PGDHi treatment reverses this regenerative deficit, enhancing MuSC proliferation and promoting myogenic differentiation into new myofibers. GLP-1 receptor agonism has been linked to lean mass loss and blunted muscle growth programs. The ability of PGDHi to restore regenerative capacity highlights a synergistic benefit of a combined drug treatment: it preserves the metabolic advantages of GLP-1RA therapy while maintaining the muscle’s ability to repair and regrow after injury.

## Discussion

GLP-1 receptor agonists have emerged as transformative therapies for obesity, leading to substantial and sustained weight loss across diverse populations (45). However, a portion of this weight loss is attributable to reductions in lean mass, including skeletal muscle (7, 15, 23, 24, 46). There is growing concern that weight loss may come at the expense of strength, which is essential for mobility and quality of life (15, 46). Indeed, since many individuals discontinue GLP-1RA therapy (47, 48), there is a risk that fat mass will be regained after treatment cessation while muscle mass will not be restored (49), increasing the likelihood of an obese frail population. Therefore, strategies that preserve or augment muscle during pharmacologic weight loss are of particular interest. Here, we show that 15-PGDH inhibition overcomes the muscle regenerative deficit observed with GLP-1RA– induced weight loss, by enhancing muscle stem cell function and increasing muscle strength.

Despite broad consensus that semaglutide reduces skeletal muscle mass in obese mice (17, 31, 50) and humans (7, 51, 52), its impact on muscle strength is less clear, with results varying by species and by the functional assay used. In mice, some studies report impaired muscle function using forelimb grip strength (50), or ex vivo twitch force (31), whereas other clinical studies report little to no change in physical performance or strength assessed by hand grip strength (53) and chair rise and gait speed (32). Here, we extend prior work by measuring in vivo maximal tetanic force, a physiologically relevant endpoint that integrates neural activation with whole-muscle force production and is less confounded by motivation, learning, or limb use than voluntary tests such as grip strength. Notably, we find that with semaglutide treatment the reduction in muscle mass is not accompanied by a decrement in strength under steady-state conditions, as maximal tetanic force and specific force were not affected. Furthermore, co-treatment of semaglutide and a 15-PGDH inhibitor yielded similar results to semaglutide alone, indicating that 15-PGDH inhibition does not interfere with the weight loss effects of semaglutide, while maintaining muscle function.

With aging and with weight loss there is a concern that muscle strength will be lost and therefore building muscle through exercise is strongly advised. We reasoned that at steady state deficits may not be detected that would become apparent upon anabolic challenge. We therefore tested whether semaglutide alters MuSC-dependent regenerative capacity and whether PGDHi can restore this muscle-building response. Satellite cells are dedicated muscle stem cells (MuSC) that are juxtaposed to myofibers in a quiescent state, poised to spring into action when damage occurs. They are essential to skeletal muscle building in response to physiological challenges such as exercise (54–56) or trauma (57, 58). Because GLP-1 receptor agonist–induced weight loss is accompanied by a reduction in lean mass triggered by low calorie intake and high catabolism, we sought to determine whether MuSC function is altered in obese mice undergoing semaglutide-induced weight loss and if so whether 15-PGDH inhibition could surmount this deficit. To test this, as a proxy for muscle growth as occurs in exercise, we used notexin injury as an in vivo assay for myogenic repair capacity that maximally challenges MuSC activation, expansion and fusion to rebuild myofibers. We find that semaglutide blunts muscle fiber regeneration, but that this impairment is overcome by 15-PGDH inhibition. Semaglutide-treated obese mice displayed significantly smaller regenerating myofibers, indicative of reduced myofiber growth. This phenotype resembles that observed in fasting models, where energy deprivation blunts regeneration and reduces fiber size compared to ad libitum-fed controls (59). Mechanistically, caloric restriction has been shown to impede regeneration by driving MuSCs into a deeper quiescent state, delaying their cell cycle entry (59). This is consistent with semaglutide’s effects on appetite suppression. Consistent with this finding, we show that the reduced MuSC proliferation under nutrient-depleted culture conditions is partially rescued by PGE2, in good agreement with the known role of PGE2 in enhancing MuSC function (our refs).

We have previously shown that MuSCs require PGE2 to function. In the absence of this signaling pathway, they and fail to proliferate and frequently die. Following injury, force is lost due to the inability of MuSCs to respond to PGE2 in the absence of the EP4 receptor or in the presence of an NSAID that blocks endogenous PGE2 synthesis. Moreover, the proliferative deficit of aged MuSCs is overcome upon exposure to PGE2 ex vivo as is evident by their superior engraftment following transplantation. Strikingly, the response of MuSCs in the muscles of aged MuSCs in situ to a single exposure of PGE2 post injury is so profound that muscle mass and strength are significantly increased two weeks later. (20, 22). Furthermore, inhibition of 15-PGDH, the major enzyme that degrades PGE2, elevates PGE2 levels and not only acts to enhance stem cell function, but also muscle fiber mitochondrial and contractile function and neuromuscular connectivity in aged mice (19, 21). Importantly, 15-PGDH inhibition allows physiological modulation of PGE2, to pre-injury or young levels. Here we extend these findings by showing that 15-PGDH inhibition overcomes a regenerative deficit produced by semaglutide.

To our knowledge it has not previously been reported that in obese mice regeneration of muscle after damage leads to an increase in calcified regions in obese mice. Such calcified regions interfere with efficacious contractile function (60). We found that regenerating muscles of vehicle-treated obese mice had large areas of calcification, a hallmark of pathological remodeling seen in muscle-wasting diseases such as Duchenne muscular dystrophy (DMD), a disease of stem cell exhaustion and regenerative failure. This calcification may arise from aberrant TGF-β signaling by fibroadipogenic progenitors (FAPs) (61, 62), and is indicative of failed or misdirected regeneration. Notably, semaglutide treatment prevented the accumulation of calcified lesions, likely because it reduces circulating TGF-β which is upregulated in obesity (63), highlighting a protective effect of GLP-1 RA against maladaptive muscle tissue remodeling. However, this benefit comes at the cost of reduced myofiber growth. Importantly, the 15-PGDH inhibitor (PGDHi) synergizes with semaglutide in obese mice to restore fiber size and increase force.

Several therapeutic strategies aimed at enhancing muscle mass in the context of GLP-1RA mediated weight loss have focused on inhibiting the myostatin/activin signaling pathway (17, 64), given its well-characterized role in muscle wasting. While these approaches have demonstrated substantial hypertrophic effects in preclinical models, translation to human populations has shown mixed results. Clinical trials with myostatin inhibitors alone have reported increases in lean mass without corresponding gains in muscle strength (65, 66). More recently, Phase 2 combination trials pairing GLP1-RA therapies with myostatin/ActRII-pathway inhibitors (e.g., Regeneron and Scholar Rock programs) have reported improved body-composition and preservation of lean mass, but whether these regimens affect force remains to be determined in mice and humans.

In conclusion, our results identify 15-PGDH inhibition as a therapeutic strategy with the potential to overcome the muscle regenerative deficits associated with semaglutide-induced weight loss without blunting semaglutide’s beneficial effects on body weight reduction. These findings provide mechanistic evidence for a synergistic co-therapy, that will enable robust fat loss while preserving muscle regenerative capacity and strength.

## Methods

### Mouse studies

Animal experiments were performed in accordance with procedures approved by the Institutional Animal Care and Use Committee of the Stanford Animal Care and Use Committee (APLAC) protocol number #ALPAC-32982. C57BL/6J male mice were purchased from the Jackson Laboratory and were maintained on a high-fat diet (HFD, 60% kcal from fat, # D12492, Research Diets) for 12 weeks. All mice were in good health and housed in a temperature-controlled (20-22°C) room on a 12-hour light/dark cycle with ad-lib access to food and water. Mice weights were daily recorded. At the end of the experiments, body and tissue weights were recorded. Tissues were collected and frozen for further analysis.

### GLP-1 RA and PGDHi administration

Semaglutide was used as GLP-1 RA at a concentration of 120 μg/kg and SW033291 (ApexBio cat #A8709; named PGDHi in this study) was used at a concentration of 5 mg/kg. Mice were treated with both drugs or the corresponding vehicle controls daily via intraperitoneal injections. Semaglutide was diluted in 90% saline solution, 5% Kolliphor and 5% DMSO, and PGDHi was diluted in 10% ethanol, 5% Cremophor EL (Sigma-Aldrich cat # C5135), 85% D5W (Dextrose 5% Water) as previously described (26).

### Glucose tolerance tests

For glucose tolerance testing, mice were fasted for 4 hours and then received an intraperitoneal injection of glucose (1.5 g/kg body weight). Blood glucose levels were recorded at 0, 15, 30, 45, 60, 90, and 120 minutes post-injection.

### Muscle injury

We induced acute muscle injury by intramuscular injection of notexin into the gastrocnemius (GA) muscle (40 µl; 10 µg ml^−1^; Latoxan, catalog# L8104). Notexin is a snake venom–derived phospholipase A_2_ toxin that, when injected into muscle, causes rapid myofiber necrosis (67). The resulting tissue damage initiates a regenerative program characterized by activation and proliferative expansion of muscle stem cells to restore the injured muscle. Tissues were collected at 5 and 14 days post injury.

### Histology and tissue staining

After collection, tissues were embedded in O.C.T. compound and snap frozen in liquid nitrogen. Using a cryostat 10-to-15-µm sections were cut. Samples were kept at -20 °C until histochemical analysis.

For histochemical analysis, sections were fixed in 4% PFA for 10 min and washed 3 times with PBS for 10 minutes each wash. After washing, the sections were blocked with Blocking One reagent for 45 minutes, and incubated overnight at 4 °C with the corresponding first antibodies. The next day, samples were washed 3 times for 10 min with PBS-T and incubated at room temperature for one hour with the corresponding antibodies. After incubation with secondary antibodies, samples were washed 3 times with PBS for 5 min each time, and mounted in Fluoromont-G (SouthernBiotech).

For hematoxylin and eosin (H&E) staining, iWAT, BAT and livers were formalin-fixed, paraffin-embedded, and sectioned at 6 µm. Sections were deparaffinized and dehydrated with xylene and ethanol. Briefly, slides were stained with hematoxylin, washed with water and 95% ethanol, and stained with eosin for 30 minutes. Sections were then incubated with ethanol and xylene and mounted with mounting medium. Digital images were captured with an Aperio AT2 (Leica) and QuPath software was used to extract the images

### Image analysis

Stained sections were imaged using a Keyence microscope and analyzed with the BZ-X analyzer software (Keyence, Osaka, Japan). For quantifications, whole tissue cross sections were analyzed.

### Force measurements

Peak isometric torque of the ankle plantar flexors was measured as previously described (19, 21, 68, 69). Briefly, anesthetized mice (3% isoflurane mixed with oxygen) were positioned with the foot secured to a footplate connected to a servomotor (Model 300C-LR, Aurora Scientific). Percutaneous Pt-Ir electrode needles (Aurora Scientific) were inserted over the tibial nerve near the posterior-medial knee. The ankle was fixed at a 90° angle, and maximal isometric torque was elicited by stimulating the tibial nerve at 150 Hz using 0.1 ms square wave pulses. Three tetanic contractions were recorded per muscle, with one-minute intervals between trials. Measurements were performed blinded to treatment groups. For relative force assessment, values were recorded both before and one month after treatment, and percent change was calculated. Data acquisition and analysis were performed using Dynamic Muscle Data Acquisition and Analysis Software (Aurora Scientific). Force was measured longitudinally in mice undergoing semaglutide and/or PGDHi treatment at baseline, 3 weeks and 5 weeks. Specific force was calculated by normalizing plantar flexion maximal contraction to the harvested gastrocnemius muscles.

### Mouse muscle stem cells isolation

Muscle stem cells (MuSCs) were isolated from mouse hindlimb muscles using a combination of enzymatic digestion, magnetic depletion, and fluorescence-activated cell sorting (FACS). Briefly, dissected muscles were minced and digested with 0.2% collagenase type II (Worthington) for 60 min, followed by an additional 30 min digestion in collagenase/dispase (0.04 U/mL; Thermo Fisher Scientific). The resulting suspension was triturated through an 18-gauge needle to release mononuclear cells. Cells were pelleted by centrifugation at 300 g for 5 min, resuspended in FACS buffer (PBS, 2% FBS, 2 mM EDTA), and filtered through a 70 μm strainer.

For immunolabeling, cells were incubated on ice for 45 min in the dark with the following antibodies: CD11b (1:800), CD45 (1:500), Sca1 (1:200), CD31 (1:200), and α7-integrin– PE (1:200). After washing in FACS buffer, samples were incubated with streptavidin-coated magnetic beads (1:200; Miltenyi Biotec) to deplete lineage-positive cells (CD11b^+^, CD45^+^, Sca1^+^, CD31^+^). Lineage-negative cells were then enriched using magnetic activated cell sorting (MACS). The enriched fraction was subsequently analyzed by flow cytometry, and MuSCs were defined and sorted based on α7-integrin–PE positivity, using unstained controls to establish gating. This strategy reliably yielded α7-integrin^+^CD11b^−^ CD45^−^Sca1^−^CD31^−^ MuSCs with high purity for downstream culture and functional assays.

### Mouse muscle stem cells culture and treatments

Isolated MuSCs were cultured in either nutrient-depleted or regular growth medium. The nutrient-depleted medium consisted of DMEM low glucose (1 g/L) supplemented with 1 mM sodium pyruvate, 2% charcoal-stripped FBS, 5 ng/mL FGF2 (Peprotech), 1× penicillin/streptomycin, and 1.0 mM GlutaMAX (Thermo Fisher Scientific). This formulation excluded glutamine and phenol red to reduce extrinsic nutrient and signaling inputs. The regular growth medium consisted of standard DMEM supplemented with 20% FBS, 5 ng/mL FGF2, and 1× penicillin/streptomycin.

Cells were plated in 96-well plates and cultured for 48 h under the indicated conditions. Pharmacological treatments were applied as follows: PGE2 (10 μM), EP2/EP4 receptor inhibitor (10 μM), or CREB inhibitor (1 μM). For proliferation assays, cells were pulsed with EdU (5 μM) during the final 12 h of culture. At the end of the treatment period, cells were fixed and processed for EdU incorporation analysis according to the manufacturer’s protocol (Thermo Fisher Scientific).

### Statistical analyses

Data are presented as mean ± SEM and as individual data points. Statistical analyses were conducted using GraphPad Prism. Replicates and sample sizes (N), representing individual animals, are also provided in the figure legends. Prior to treatment, mice were randomized based on body weight and baseline force measurements. Group differences were analyzed using ANOVA followed by Tukey’s post hoc test for multiple comparisons. A p-value < 0.05 was considered statistically significant.

## Acknowledgments

This study was supported by the Baxter Foundation, the Milky Way Research Foundation (grant 216064) (to H.M.B.), a Stanford Cardiovascular Institute (CVI) Seed Grant award (to H.M.B.and M.N.), and the Stanford Bio-X Summer Undergraduate Research Program (awarded to I.K.). H.M.B. was also supported by NIH grants 5R01AG02096115, 1R01AG069858-01, and RHG009674A. K.J.S. was supported by R01DK125260, P30DK116074, and the American Heart Association (23IPA1042031). This research was supported in part by the Arc Institute. K.J.S. is an Arc Institute Innovation Investigator and a Weill Cancer Hub West Investigator. M.Z. was supported by K99/R00 NIH Pathway to Independence Award (K99AR081618).

## Author Contributions

M.N., J.L., K.S. and H.M.B. designed research; M.N., J.L, E.L.M., P.K., K.K., Z.Z. and S.B. performed research; M.N. and J.L. analyzed data; M.N. and H.B. wrote the manuscript, and all the authors revised the manuscript.

## References

1. D. P. Guh, et al., The incidence of co-morbidities related to obesity and overweight: A systematic review and meta-analysis. BMC Public Health 9, 88 (2009).

2. C. E. Hagberg, K. L. Spalding, White adipocyte dysfunction and obesity-associated pathologies in humans. Nat Rev Mol Cell Biol 25, 270–289 (2024).

3. B. M. Spiegelman, J. S. Flier, Obesity and the Regulation of Energy Balance. Cell 104, 531–543 (2001).

4. M. Ng, et al., Global, regional, and national prevalence of overweight and obesity in children and adults during 1980–2013: a systematic analysis for the Global Burden of Disease Study 2013. The Lancet 384, 766–781 (2014).

5. X. Zhao, et al., GLP-1 Receptor Agonists: Beyond Their Pancreatic E]ects. Front. Endocrinol. 12 (2021).

6. D. J. Drucker, Mechanisms of Action and Therapeutic Application of Glucagon-like Peptide-1. Cell Metabolism 27, 740–756 (2018).

7. J. P. H. Wilding, et al., Once-Weekly Semaglutide in Adults with Overweight or Obesity. New England Journal of Medicine (2021). 10.1056/NEJMoa2032183.

8. V. Perkovic, et al., Effects of Semaglutide on Chronic Kidney Disease in Patients with Type 2 Diabetes. New England Journal of Medicine (2024).

9. M. N. Kosiborod, et al., Semaglutide in Patients with Obesity-Related Heart Failure and Type 2 Diabetes. New England Journal of Medicine (2024). 10.1056/NEJMoa2313917.

10. L. Sjöström, et al., Effects of Bariatric Surgery on Mortality in Swedish Obese Subjects. New England Journal of Medicine 357, 741–752 (2007).

11. D. Weghuber, et al., Once-Weekly Semaglutide in Adolescents with Obesity. New England Journal of Medicine (2022). 10.1056/NEJMoa2208601.

12. M. N. Kosiborod, et al., Semaglutide in Patients with Heart Failure with Preserved Ejection Fraction and Obesity. New England Journal of Medicine 389, 1069–1084 (2023).

13. N. Sattar, et al., Cardiovascular, mortality, and kidney outcomes with GLP-1 receptor agonists in patients with type 2 diabetes: a systematic review and meta-analysis of randomised trials. The Lancet Diabetes & Endocrinology 9, 653–662 (2021).

14. Muscle matters: the effects of medically induced weight loss on skeletal muscle-The Lancet Diabetes & Endocrinology. Available at: https://www.thelancet.com/journals/landia/article/PIIS2213-8587(24)00272-9/fulltext [ Accessed 25 June 2025].

15. C. Arnold, After obesity drugs’ success companies rush to preserve skeletal muscle. Nature Biotechnology 42, 351–353 (2024).

16. R. H. Choi, et al., Semaglutide-induced weight loss improves mitochondrial energy efficiency in skeletal muscle. Obesity 33, 974–985 (2025).

17. E. Nunn, et al., Antibody blockade of activin type II receptors preserves skeletal muscle mass and enhances fat loss during GLP-1 receptor agonism. Mol Metab 80, 101880 (2024).

18. J. Chen, R. Zhou, Y. Feng, L. Cheng, Molecular mechanisms of exercise contributing to tissue regeneration. Sig Transduct Target Ther 7, 383 (2022).

19. A. R. Palla, et al., Inhibition of prostaglandin-degrading enzyme 15-PGDH rejuvenates aged muscle mass and strength. Science (2021). 10.1126/science.abc8059.

20. Y. X. Wang, et al., Multiomic profiling reveals that prostaglandin E2 reverses aged muscle stem cell dysfunction, leading to increased regeneration and strength. Cell Stem Cell (2025). 10.1016/j.stem.2025.05.012.

21. M. A. Bakooshli, et al., Regeneration of neuromuscular synapses after acute and chronic denervation by inhibiting the gerozyme 15-prostaglandin dehydrogenase. Sci Transl Med 15, eadg1485 (2023).

22. A. T. V. Ho, et al., Prostaglandin E2 is essential for efficacious skeletal muscle stem-cell function, augmenting regeneration and strength. Proc Natl Acad Sci U S A 114, 6675–6684 (2017).

23. J. Linge, A. L. Birkenfeld, I. J. Neeland, Muscle Mass and Glucagon-Like Peptide-1 Receptor Agonists: Adaptive or Maladaptive Response to Weight Loss? Circulation 150, 1288–1298 (2024).

24. C. Conte, K. D. Hall, S. Klein, Is Weight Loss–Induced Muscle Mass Loss Clinically Relevant? JAMA 332, 9–10 (2024).

25. A. R. Palla, et al., Inhibition of prostaglandin-degrading enzyme 15-PGDH rejuvenates aged muscle mass and strength. Science 371, eabc8059 (2021).

26. Y. Zhang, et al., Inhibition of the prostaglandin-degrading enzyme 15-PGDH potentiates tissue regeneration. Science 348, aaa2340 (2015).

27. C. Quarta, et al., GLP-1-mediated delivery of tesaglitazar improves obesity and glucose metabolism in male mice. Nat Metab 4, 1071–1083 (2022).

28. M. D. Martens, et al., Semaglutide Reduces Cardiomyocyte Size and Cardiac Mass in Lean and Obese Mice. JACC: Basic to Translational Science 9, 1429– 1431 (2024).

29. J. Xiang, L. Qin, J. Zhong, N. Xia, Y. Liang, GLP-1RA Liraglutide and Semaglutide Improves Obesity-Induced Muscle Atrophy via SIRT1 Pathway. Diabetes, Metabolic Syndrome and Obesity: Targets and Therapy 16, 2433–2446 (2023).

30. X. Pan, L. Yue, J. Ban, L. Ren, S. Chen, Effects of Semaglutide on Cardiac Protein Expression and Cardiac Function of Obese Mice. JIR 15, 6409–6425 (2022).

31. T. Karasawa, et al., Unexpected effects of semaglutide on skeletal muscle mass and force-generating capacity in mice. Cell Metabolism 37, 1619–1620 (2025).

32. G. L. Ditzenberger, et al., Effects of Semaglutide on Muscle Structure and Function in the SLIM LIVER Study. Clinical Infectious Diseases 80, 389–396 (2025).

33. J. McKendry, T. Stokes, J. C. Mcleod, S. M. Phillips, “Resistance Exercise, Aging, Disuse, and Muscle Protein Metabolism” in Comprehensive Physiology, (John Wiley & Sons, Ltd, 2021), pp. 2249–2278.

34. H. M. Blau, Regulating the myogenic regulators. Symp Soc Exp Biol 46, 9–18 (1992).

35. H. M. Blau, B. D. Cosgrove, A. T. V. Ho, The central role of muscle stem cells in regenerative failure with aging. Nat Med 21, 854–862 (2015).

36. A. T. V. Ho, et al., Prostaglandin E2 is essential for efficacious skeletal muscle stem-cell function, augmenting regeneration and strength. Proc Natl Acad Sci U S A 114, 6675–6684 (2017).

37. X. Fu, et al., Obesity Impairs Skeletal Muscle Regeneration Through Inhibition of AMPK. Diabetes 65, 188–200 (2015).

38. L. G. M. Heezen, et al., Spatial transcriptomics reveal markers of histopathological changes in Duchenne muscular dystrophy mouse models. Nat Commun 14, 4909 (2023).

39. D. W. Hammers, et al., The D2.mdx mouse as a preclinical model of the skeletal muscle pathology associated with Duchenne muscular dystrophy. Sci Rep 10, 14070 (2020).

40. D. A. G. Mázala, et al., Altered muscle niche contributes to myogenic deficit in the D2-mdx model of severe DMD. Cell Death Discov. 9, 224 (2023).

41. B. D. Cosgrove, et al., Rejuvenation of the muscle stem cell population restores strength to injured aged muscles. Nat Med 20, 255–264 (2014).

42. L. Walter, B. Cosgrove, Transcriptomic analysis of skeletal muscle regeneration across mouse lifespan identifies altered stem cell states. Dryad. 10.5061/DRYAD.KKWH70SBV. Deposited 27 November 2024.

43. D. W. McKellar, et al., Large-scale integration of single-cell transcriptomic data captures transitional progenitor states in mouse skeletal muscle regeneration. Commun Biol 4, 1–12 (2021).

44. B. D. Cosgrove, et al., Rejuvenation of the muscle stem cell population restores strength to injured aged muscles. Nat Med 20, 255–264 (2014).

45. Z. Zheng, et al., Glucagon-like peptide-1 receptor: mechanisms and advances in therapy. Sig Transduct Target Ther 9, 234 (2024).

46. C. M. Prado, S. M. Phillips, M. C. Gonzalez, S. B. Heymsfield, Muscle matters: the effects of medically induced weight loss on skeletal muscle. The Lancet Diabetes & Endocrinology 12, 785–787 (2024).

47. P. P. Gleason, et al., Real-world persistence and adherence to glucagon-like peptide-1 receptor agonists among obese commercially insured adults without diabetes. JMCP 30, 860–867 (2024).

48. P. J. Rodriguez, et al., Discontinuation and Reinitiation of Dual-Labeled GLP-1 Receptor Agonists Among US Adults With Overweight or Obesity. JAMA Netw Open 8, e2457349 (2025).

49. J. P. H. Wilding, et al., Weight regain and cardiometabolic effects after withdrawal of semaglutide: The STEP 1 trial extension. Diabetes, Obesity and Metabolism 24, 1553–1564 (2022).

50. S. Jeromson, et al., Semaglutide impacts skeletal muscle to a similar extent as caloric restriction in mice with diet-induced obesity. The Journal of Physiology n/a.

51. K. Prokopidis, Glucagon-like peptide-1 receptor agonists and muscle strength changes in older adults: Risks beyond muscle mass reductions. British Journal of Pharmacology n/a.

52. M. Alissou, et al., Impact of Semaglutide on fat mass, lean mass and muscle function in patients with obesity: The SEMALEAN study. Diabetes, Obesity and Metabolism 28, 112–121 (2026).

53. S. Volpe, et al., Once-Weekly Semaglutide Induces an Early Improvement in Body Composition in Patients with Type 2 Diabetes: A 26-Week Prospective Real-Life Study. Nutrients 14 (2022).

54. D. A. Englund, et al., Depletion of resident muscle stem cells negatively impacts running volume, physical function, and muscle ?ber hypertrophy in response to lifelong physical activity. American Journal of Physiology-Cell Physiology 318, C1178–C1188 (2020).

55. A. P. Sharples, D. C. Turner, Skeletal muscle memory. American Journal of Physiology-Cell Physiology 324, C1274–C1294 (2023).

56. T. P. Saliu, et al., Satellite cell dynamics during skeletal muscle hypertrophy. Biochem Soc Trans 52, 1921–1926 (2024).

57. S. M. Hindi, D. P. Millay, All for One and One for All: Regenerating Skeletal Muscle. Cold Spring Harb Perspect Biol 14, a040824 (2022).

58. F. Relaix, et al., Perspectives on skeletal muscle stem cells. Nat Commun 12, 692 (2021).

59. D. I. Benjamin, et al., Fasting induces a highly resilient deep quiescent state in muscle stem cells via ketone body signaling. Cell Metabolism 34, 902-918.e6 (2022).

60. N. A. Mignemi, et al., Plasmin Prevents Dystrophic Calcification After Muscle Injury. J Bone Miner Res 32, 294–308 (2017).

61. D. A. Mázala, et al., TGF-β-driven muscle degeneration and failed regeneration underlie disease onset in a DMD mouse model. JCI Insight 5, e135703. 135703 (2020).

62. J. B. Lees-Shepard, et al., Activin-dependent signaling in ?bro/adipogenic progenitors causes fibrodysplasia ossificans progressiva. Nat Commun 9, 471 (2018).

63. H. Yadav, et al., Protection from Obesity and Diabetes by Blockade of TGF-β/Smad3 Signaling. Cell Metabolism 14, 67–79 (2011).

64. J. W. Mastaitis, et al., GDF8 and activin A blockade protects against GLP-1– induced muscle loss while enhancing fat loss in obese male mice and non-human primates. Nat Commun 16, 4377 (2025).

65. A. A. Amato, et al., Treatment of sporadic inclusion body myositis with bimagrumab. Neurology 83, 2239–2246 (2014).

66. K. R. Wagner, et al., A phase I/IItrial of MYO-029 in adult subjects with muscular dystrophy. Annals of Neurology 63, 561–571 (2008).

67. L. Boldrin, A. Neal, P. S. Zammit, F. Muntoni, J. E. Morgan, Donor Satellite Cell Engraftment is Significantly Augmented When the Host Niche is Preserved and Endogenous Satellite Cells are Incapacitated. Stem Cells 30, 1971–1984 (2012).

68. K. A. Sheth, et al., Muscle strength and size are associated with motor unit connectivity in aged mice. Neurobiol Aging 67, 128–136 (2018).

69. E. L. Mintz, J. A. Passipieri, D. Y. Lovell, G. J. Christ, Applications of In Vivo Functional Testing of the Rat Tibialis Anterior for Evaluating Tissue Engineered Skeletal Muscle Repair. J Vis Exp 54487 (2016). 10.3791/54487.

