## Supplemental Figures for "15-PGDH Inhibition Overcomes Muscle Regenerative Deficit Seen With GLP1-Receptor Agonist–Induced Weight Loss"

### Figure S1

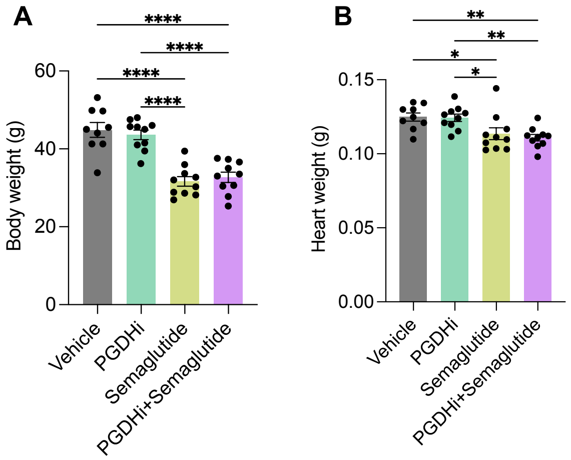


**Fig. S1. Effect of semaglutide and/or PGDHi on body and heart weights in obese mice**
(A) Body weight in grams at the end of the treatment.
(C) Wet heart tissue weight.
n=8-10 animals per group. Data are presented as mean ± S.E.M. Statistic significance was analyzed by two-way ANOVA for multiple comparisons. * $p < 0.05$; ** $p < 0.01$; *** $p < 0.001$; **** $p < 0.0001$.

### Figure S2

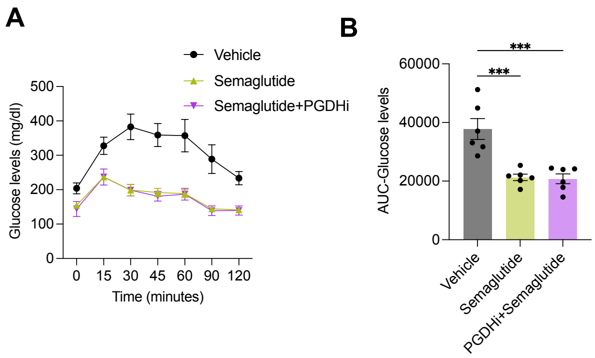


**Fig. S2. Semaglutide improves glucose tolerance in obese mice.**
(A) Blood glucose tolerance test.
(B) Area under the curve (A.U.C) of the glucose tolerance test.
(n=8-10 animals per group. Data are presented as mean ± S.E.M. Statistic significance was analyzed by two-way ANOVA for multiple comparisons. * $p < 0.05$; ** $p < 0.01$; *** $p < 0.001$; **** $p < 0.0001$.

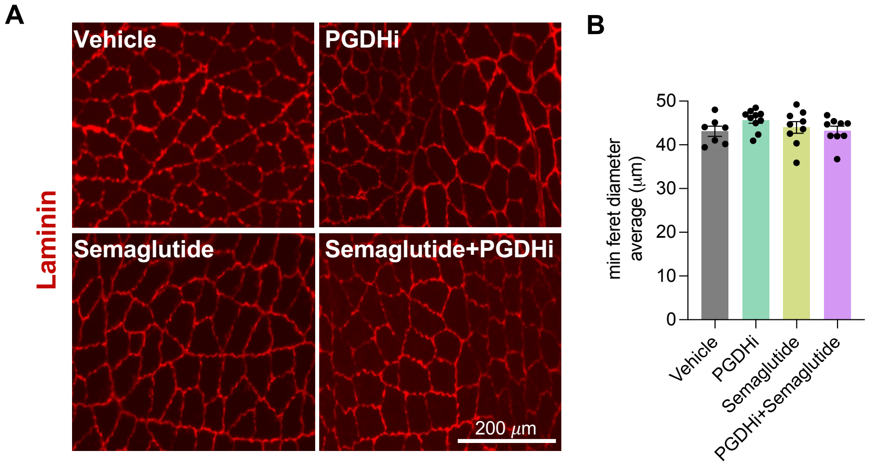


**Fig. S3. Semaglutide and/or PGDHi do not affects myofibers size in obese mice.**
(A) Representative images of gastrocnemius cross sections at 5 weeks post treatment. Laminin, Red.
(B) Quantification of fibers minimum Feret diameter.
n=8-10 animals per group. Data are presented as mean ± S.E.M. Statistic significance was analyzed by two-way ANOVA for multiple comparisons.

### Figure S4

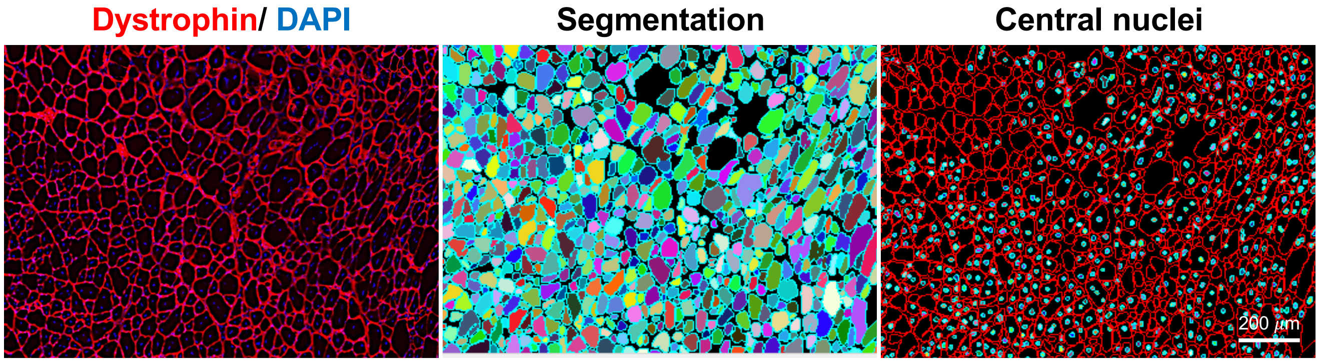


**Figure S4.** Representative image depicting the fiber segmentation and quantification method for regenerating fibers (central nuclei fibers) fiber size. Images were acquired by Keyence microscope and analyzed using BZ-X analyzer software (Keyence, Osaka, Japan).

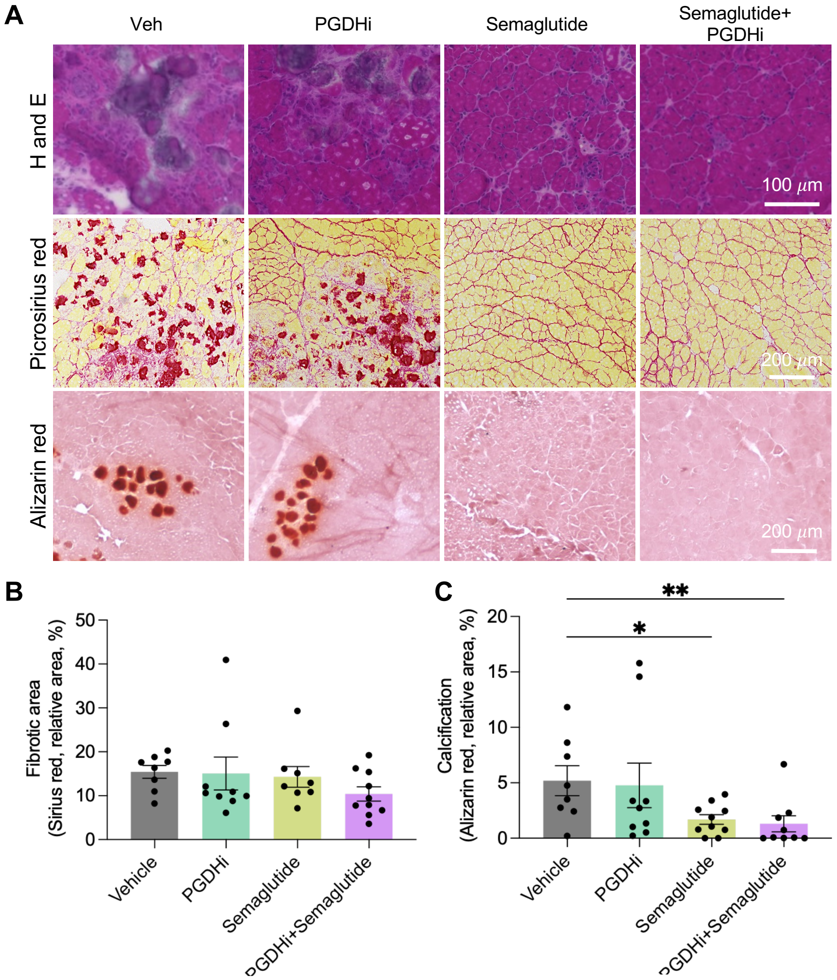


**Figure S5. Co-administration of PGDHi and semaglutide counters obesity-associated impaired regeneration.**

(A) Representative images of gastrocnemius muscles at 2 weeks post injury. From top to bottom: H&E staining; Picrosirius red staining; and Alizarin red staining.

(B) Quantification of Fibrotic area from Picrosirius red staining.

(C) Quantification of calcified area from Alizarin red staining.

n=8-10 animals per group. Data are presented as mean ± S.E.M. Statistic significance was analyzed by two-way ANOVA for multiple comparisons. * $p < 0.05$; ** $p < 0.01$; *** $p < 0.001$; **** $p < 0.0001$.

### Figure S6

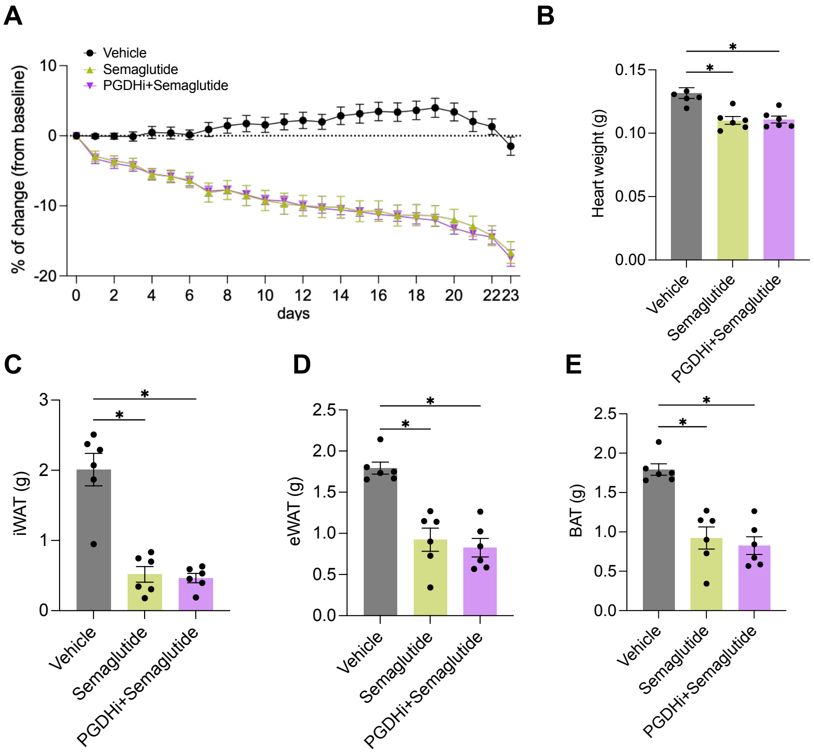


**Figure S6. Body weight and tissue weight postinjury in obese mice.**

(A) Body weight change in grams.

(B) Wet tissues weight. for the heart.

(C) Wet tissue weights for iWAT (inguinal adipose tissue), eWAT (epididymal adipose tissue), and BAT (brown adipose tissue).

n=6 animals per group. Data are presented as mean ± S.E.M. Statistic significance was analyzed by two-way ANOVA for multiple comparisons. * $p < 0.05$; ** $p < 0.01$; *** $p < 0.001$; **** $p < 0.0001$.
